# TGF-β drives the conversion of conventional NK cells into uterine tissue-resident NK cells to support murine pregnancy

**DOI:** 10.1101/2025.11.18.688992

**Authors:** Josselyn D. Barahona, Liping Yang, D. Michael Nelson, Wayne M. Yokoyama

## Abstract

Tissue microenvironments shape lymphocyte differentiation to align immune function with local physiological demands. Uterine natural killer cells are critical for reproductive success, yet the molecular cues in the uterus that instruct their specialized identities remain incompletely understood. Here, we identify a TGF-β–dependent differentiation pathway by which circulating conventional NK cells convert into uterine tissue-resident NK cells during murine pregnancy. Loss of TGF-β receptor II expression in *Ncr1*-expressing cells disrupted this conversion, markedly reducing tissue-resident NK cells in the gravid uterus. Impaired TGF-β–driven uterine tissue-resident NK cell differentiation during murine pregnancy led to abnormal spiral artery remodeling and increased fetal resorption rates at midgestation, ultimately reducing litter sizes at birth. Collectively, these findings define TGF-β as a pivotal driver of tissue-resident NK cell differentiation in the gravid uterus and establish a mechanistic framework through which the uterine microenvironment programs NK cell identity to meet the physiological demands of gestation.

## Introduction

Lymphocyte differentiation in response to local environmental cues is a fundamental mechanism by which the immune system adapts to tissue-specific demands. Outside of circulation, conventional NK (cNK) cells undergo regulated differentiation within tissue microenvironments that impose site-specific transcriptional programs to generate phenotypically and functionally discrete NK cell subsets^1–6^. In the uterus, this process gives rise to uterine NK (uNK) cells–a heterogeneous lymphocyte population composed primarily of tissue-resident NK (trNK) cells, with a minor subset of cNK cells–that are thought to mediate critical gestational adaptations, including placental vascular remodeling, trophoblast differentiation, and fetal development, through the secretion of cytokines and growth factors^2,7–16^. The importance of uNK cells in pregnancy is evidenced by our previous findings demonstrating that their absence results in reduced litter sizes and increased fetal resorption rates in mice^17^. In humans, disruptions in uNK cell abundance or function have also been associated with serious obstetrical disorders, including recurrent miscarriage and preeclampsia^18–23^. Despite their importance for reproductive success, the mechanisms by which uterine trNK cells differentiate and acquire their specialized functional identities within the gravid uterus remain elusive.

Recent investigations into the developmental origins of uterine trNK cells have suggested that local molecular cues govern their differentiation. Our prior work identified Eomesodermin (Eomes) as a central transcription factor driving the establishment of trNK cells in both the virgin and pregnant murine uterus, suggesting these cells arise from precursors in the cNK cell lineage^17^. In line with this lineage relationship, parabiosis studies in virgin mice demonstrated that cNK cells in the peripheral vasculature can migrate into the uterus and adopt phenotypic characteristics consistent with uterine trNK cells^24^. Parallel findings in humans further support this differentiation pathway as analyses of endometrial biopsies from uterine transplant recipients revealed that uNK cells carry the recipient genotype, indicating a blood-borne origin for human uterine trNK cells^25^. Together, these findings support a model in which uterine trNK cells arise from hematogenous cNK cells that traffic into the pregnant uterus, where local environmental cues orchestrate their terminal differentiation.

While significant progress has been made in defining the developmental origins of uterine trNK cells, the molecular factors that instruct their differentiation within uterine tissues remain incompletely understood. In the virgin uterus, transforming growth factor (TGF)-β may be a central mediator steering the differentiation of uterine trNK cells. In the virgin uterus, trNK cells depend on sustained autocrine TGF-β signaling to maintain their population, suggesting that continuous, cell-intrinsic TGF-β signaling is critical for preserving tissue-specific NK cell identities within the uterine microenvironment^26^. Corroborating single-cell transcriptomic analyses show that, at steady-state, murine uNK cells exhibit distinct transcriptional programs enriched for TGF-β response genes, suggesting that TGF-β imprints the uNK cell compartment to drive trNK cell differentiation^27^. In humans, CD16^+^ NK cells in the peripheral blood have the potential to convert into CD16^-^ NK cells that phenotypically resemble decidual NK cells following exposure to TGF-β *in vitro*, providing evidence of a conserved role for TGF-β in promoting the phenotype of trNK cells^28^. Together, these findings position TGF-β signaling as a critical driver of uterine trNK cell identity and functional specialization within the virgin uterus. Whether TGF-β mediates this differentiation during murine pregnancy, however, remains unknown.

In this study, we identified an *in vivo* TGF-β–dependent differentiation process through which circulating cNK cells give rise to uterine trNK cells in the gravid mouse uterus. This differentiation is tightly linked to the physiological adaptations of pregnancy, as disruption of TGF-β signaling in *Ncr1*-expressing cells during murine gestation impaired spiral artery remodeling and increased resorption rates at midgestation, culminating in reduced litter sizes at birth. Collectively, this work defines a mechanistic framework in which TGF-β governs the *in vivo* differentiation and functional specialization of uterine trNK cells, thereby aligning lymphocyte differentiation to the physiological requirements of pregnancy.

## Results

### Peripheral cNK cells differentiate into trNK cells in the pregnant murine uterus

The pregnant uterus is characterized by three innate lymphoid cell subsets distinguished by the expression of CD49a, CD49b, and Eomes^2,7^. CD49a^+^ Eomes^+^ trNK cells and CD49a^+^ Eomes^-^ type 1 innate lymphoid cells (ILC1s) reside within uterine tissues, whereas CD49b^+^ Eomes^+^ cNK cells circulate through the peripheral vasculature^24^. To determine whether cNK cells migrate into the pregnant uterus, we intravenously administered a fluorescently labeled anti-CD45.2 antibody *in vivo* prior to euthanasia, allowing us to discriminate circulating lymphocytes (CD45.2–PE-Cy7^+^) from those residing within uterine tissues (CD45.2–PE-Cy7^-^). CD49a^+^ Eomes^+^ trNK cells and CD49a^+^ Eomes^-^ ILC1s within implantation sites were not labeled with our circulating antibody, confirming their residency within tissues of the gravid uterus. Interestingly, intravascular labeling of circulating lymphocytes revealed a population of extravascular CD49b^+^ Eomes^+^ cNK cells within implantation sites at gestational day (gd) 6.5 that increased at gd 14.5, suggesting that peripheral cNK cells extravasate into the pregnant murine uterus (**Figure 1 A, B**).

**Figure 1.**
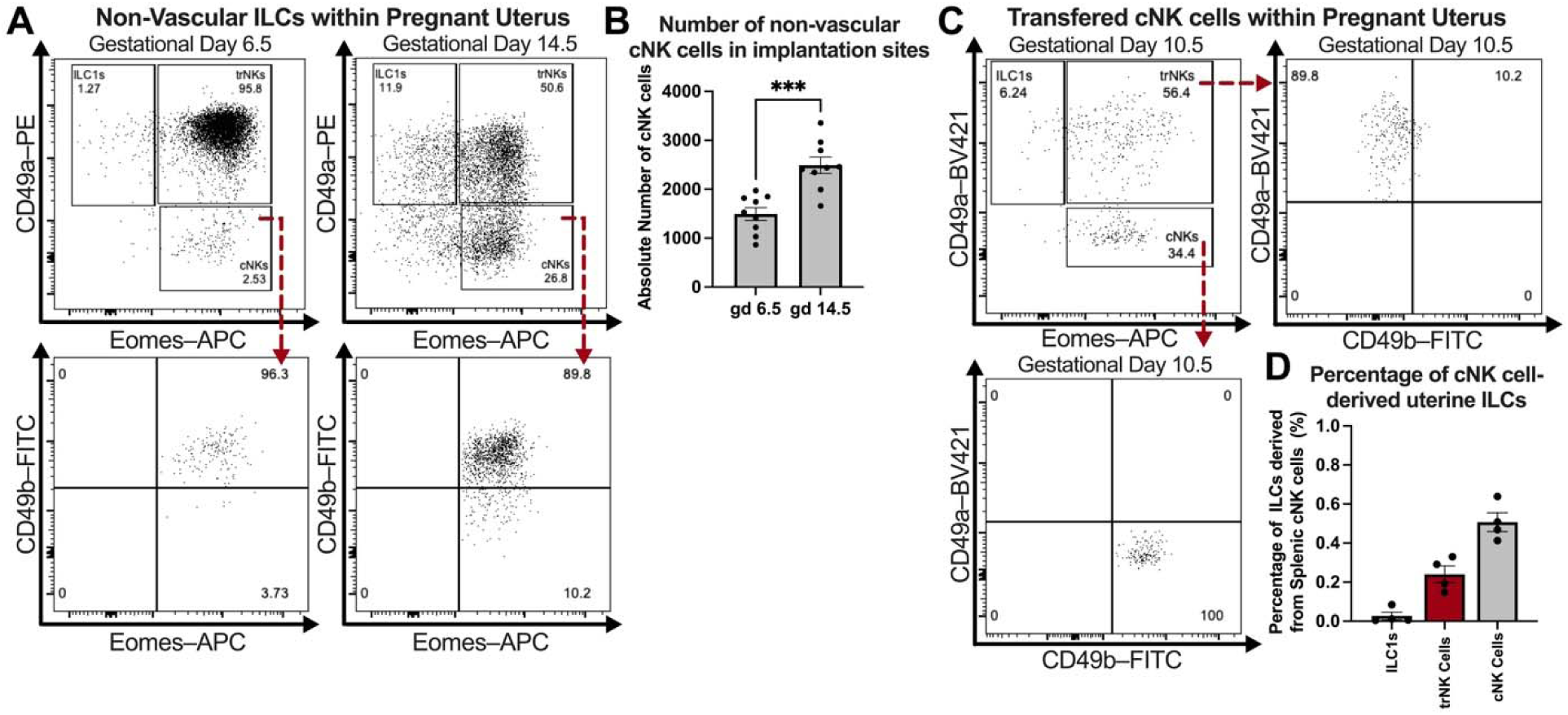
Peripheral cNK cells extravasate into the pregnant uterus and acquire a uterine trNK cells phenotype. (**A**) Representative flow plots depicting the presence of non-vascular CD49b^+^ Eomes^+^ cNK cells within the gravid uterus of wildtype mice intravascularly labeled with anti-CD45.2 antibody *in vivo* at gds 6.5 and 14.5 (gd 6.5: C57BL/6 dams, *n*=3, implantation sites *n*=9; gd 14.5: C57BL/6 dams, *n*=3, implantation sites *n*=9). Gating strategy: Live, Single Cells; CD3^-^ CD19^-^ CD45.1^-^ CD45.2–PE-Cy7^-^ CD45.2–Pacific Blue^+^ NK1.1^+^ NKp46^+^ cells. (**B**) Absolute cell counts of non-vascular CD49b^+^ Eomes^+^ cNK cells within the gravid uterus of wildtype mice at gds 6.5 and 14.5. (**C**) Concatenated flow plots of implantation sites showing that adoptively transferred cNK cells in pregnant uterus of wildtype dams upregulate CD49a and down regulate CD49b by gd 10.5, acquiring a CD49a^+^ CD49b^-^ Eomes^+^ phenotype characteristic of uterine trNK cells (C57BL/6 dams *n*=4). Here, 2.5x10^6^ CD45.2^+^ CD3^-^ CD19^-^ NK1.1^+^ NKp46^+^ CD49b^+^ splenic cNK cells were adoptively transferred into pregnant C57BL/6-CD45.1 dams at gd 0.5, and the receptor profile of these cells was subsequently assessed at gd 10.5. Gating strategy: Live, Single Cells; CD3^-^ CD19^-^ CD45.1^-^ CD45.2–PE-Cy7^-^ CD45.2–PE^+^ NK1.1^+^ NKp46^+^ cells. (**D**) Proportion of uterine ILC subsets derived from adoptively transferred splenic cNK cells in the pregnant uterus of wildtype dams. Statistics were calculated using unpaired *t tests* with the Mann-Whitney correction. Error bars indicate SEM; *** *p* < 0.001.

To assess whether peripheral cNK cells migrating into the pregnant uterus could differentiate into uterine trNK cells, we adoptively transferred CD45.2^+^ splenic cNK cells into C57BL/6 CD45.1 dams mated with C57BL/6 CD45.1 males at gd 0.5. This mating strategy allowed us to distinguish donor maternal lymphocytes from fetal lymphocytes at the maternal-fetal interface. By gd 10.5, a subset of adoptively transferred splenic cNK cells upregulated CD49a and downregulated CD49b, acquiring a phenotype characteristic of CD49a^+^ Eomes^+^ uterine trNK cells (**Figure 1 C, D**). Together, these findings reveal a previously unrecognized plasticity of peripheral cNK cells *in vivo* during murine pregnancy, enabling them to convert into uterine trNK cells within the gravid uterus.

### TGF-β signaling drives the differentiation of trNK cells in the pregnant murine uterus

Inasmuch as it has been shown that TGF-β can induce an ILC1-like phenotype that can encompass trNK cells^28–31^, we sought to examine the possible role of TGF-β in the differentiation of uNK cells. Unlike peripheral cNK cells in the spleen, we found that cNK cells extravasating into the gravid uterus upregulated expression of TGF-β receptor II, suggesting that TGF-β signaling could indeed mediate their differentiation into uterine trNK cells during gestation (**Figure 2 A**). To test this, *TGF-*β*RII^fl/fl^* mice were crossed with *Ncr1^iCre^* mice to generate mice that lack TGF-βRII on NKp46^+^ NK cells and ILC1s. To ensure we exclude a confounding effect of Ncr1^iCre^ expression, we profiled the uterine innate lymphoid compartment in pregnant Ncr1^iCre^ dams at gestational day 6.5. No differences were observed in the absolute number of trNK cells, cNK cells, or ILC1s relative to wildtype controls (**Figure S1 A-D**), and implantation site number and resorption rates were likewise unchanged (**Figure S1 E-F**). These data indicate that Ncr1^iCre^ expression alone does not perturb uterine ILC composition or early pregnancy outcomes. There was no difference noted in the total number and frequency of splenic cNK cells between TGF-βRII^Ncr1Δ^ and littermate control dams during gestation (**Figure 2 B, C**). However, in the pregnant uterus, CD49a^+^ Eomes^-^ ILC1s were markedly reduced in implantation sites of TGF-βRII^Ncr1Δ^ dams, paralleling the reduction of ILC1s previously reported in the virgin uterus of TGF-βRII^Ncr1Δ^ female mice^26^. Importantly, loss of TGF-β receptor II expression in *Ncr1*-expressing cells also significantly reduced the number of CD49a^+^ Eomes^+^ trNK cells in the gravid uterus of TGF-βRII^Ncr1Δ^ dams. This reduction of uterine trNK cells was accompanied by a small increase in the absolute number and frequency of CD49b^+^ Eomes^+^ cNK cells within the pregnant uterus of TGF-βRII^Ncr1Δ^ dams (**Figure 2 D, E**). Collectively, these findings suggest that a TGF-β–driven differentiation pathway directs the conversion of peripheral cNK cells into uterine trNK cells during murine pregnancy.

**Figure 2.**
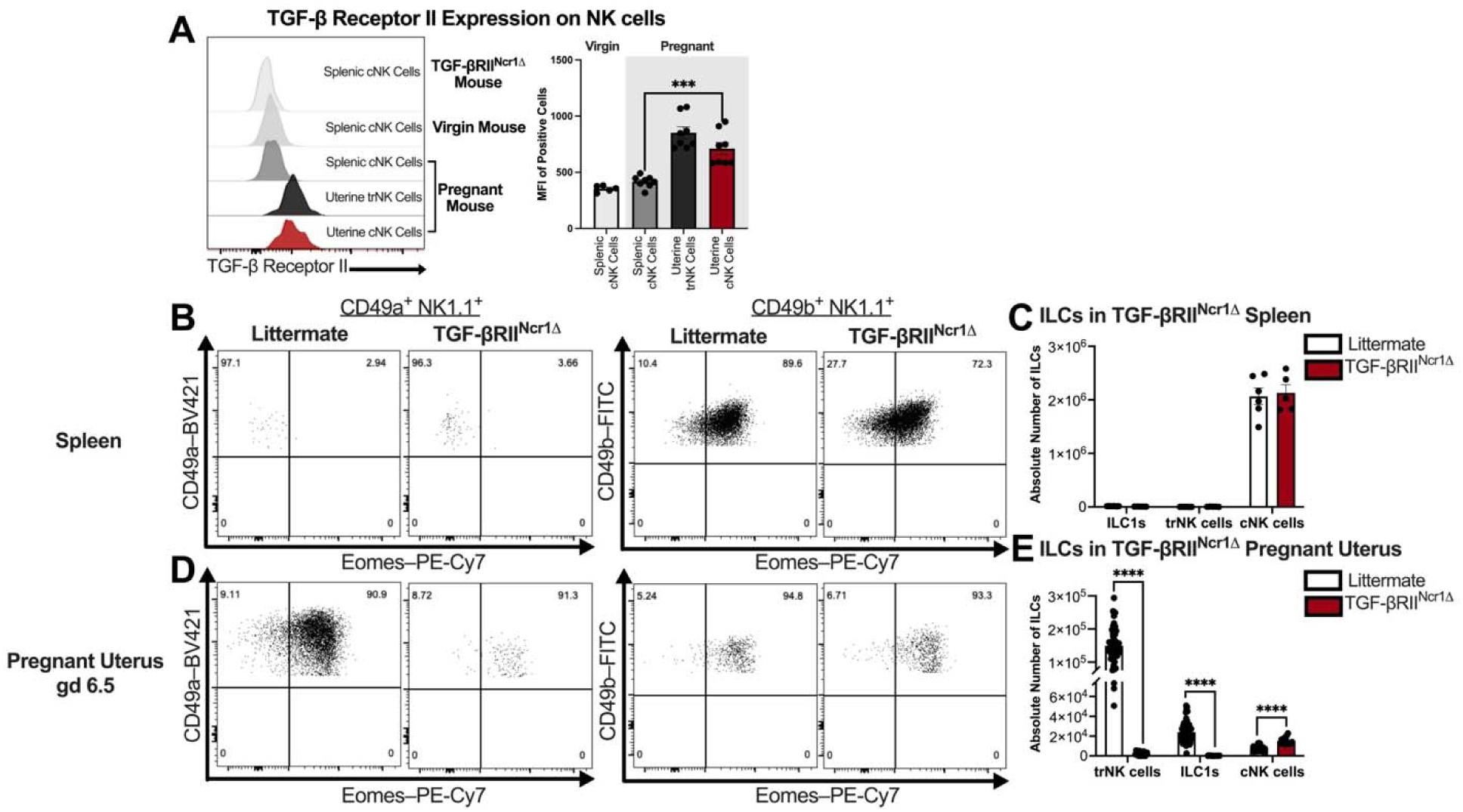
Loss of TGF-β Signaling in *Ncr1*-expressing cells impairs uterine trNK cell differentiation in pregnant mice. (**A**) Representative histograms depicting TGF-β Receptor II expression on splenic NK cells from virgin TGF-βRII^Ncr1Δ^ and wildtype mice as well as splenic and uterine NK cell subsets from pregnant wildtype mice at gd 10.5 (virgin TGF-βRII^Ncr1Δ^ mice, *n*=2; virgin mice: C57BL/6, *n*=5; gd 10.5: C57BL/6 dams, *n*=8, implantation sites *n*=8). MFI, median fluorescent intensity. Gating strategy: Live, Single Cells; CD3^-^ CD19^-^ CD45.1^-^ CD45.2^+^ NK1.1^+^ NKp46^+^ cells. (**B**) Representative flow plots showing the expression of CD49a, CD49b, and Eomes across ILC subsets in the pregnant spleens of littermate control and TGF-βRII^Ncr1Δ^ dams at gd 6.5 (Littermates, *n*=6; TGF-βRII^Ncr1Δ^, *n*=5). Gating strategy: Live, Single Cells; CD3^-^ CD19^-^CD45.1^-^ CD45.2^+^ NK1.1^+^ NKp46^+^ cells. (**C**) Absolute cell counts of CD49a^+^ Eomes^+^ trNK cells, CD49a^+^ Eomes^-^ ILC1s, and CD49b^+^ Eomes^+^ cNK cells in the spleens of pregnant littermate control and TGF-βRII^Ncr1Δ^ dams at gd 6.5. (**D**) Representative flow plots showing the expression of CD49a, CD49b, and Eomes across ILC subsets in the gravid uterus of littermate control and TGF-βRII^Ncr1Δ^ dams at gd 6.5 (Littermates, *n*=6, implantation sites *n*=54; TGF-βRII^Ncr1Δ^, *n*=5, implantation sites *n*=15). (**E**) Absolute cell counts of CD49a^+^ Eomes^+^ trNK cells, CD49a^+^ Eomes^-^ ILC1s, and CD49b^+^ Eomes^+^ cNK cells in the gravid uterus of littermate control and TGF-βRII^Ncr1Δ^ dams at gd 6.5. Statistics were calculated using unpaired *t tests* with the Mann-Whitney correction. Error bars indicate SEM; *** *p* < 0.001; and **** *p* < 0.0001.

### Impaired trNK cell differentiation in the absence of TGF-β signaling compromises pregnancy outcomes

Having established that TGF-β signaling drives the differentiation of trNK cells in the pregnant uterus, we next examined whether disrupting this pathway affects pregnancy outcomes. Litter size, pup birth weight, and gestational length at first parturition were compared between littermate control and TGF-βRII^Ncr1Δ^ dams mated with C57BL/6 CD45.1 males. TGF-βRII^Ncr1Δ^ dams had reduced litter sizes at birth compared to littermate controls (**Figure 3 A**). The birth weights of neonatal pups were not affected by the loss of TGF-β receptor II expression in *Ncr1*-expressing cells, with no relationship observed between litter size and pup birth weight (**Figure 3 B**). Furthermore, gestation proceeded over a similar timeframe in both TGF-βRII^Ncr1Δ^ dams and littermate control dams (**Figure 3 C)**. The reduction in litter size observed in TGF-βRII^Ncr1Δ^ dams suggests impaired TGF-β–dependent uterine trNK cell differentiation in the gravid murine uterus disrupts gestational adaptations crucial for fetal survival.

**Figure 3.**
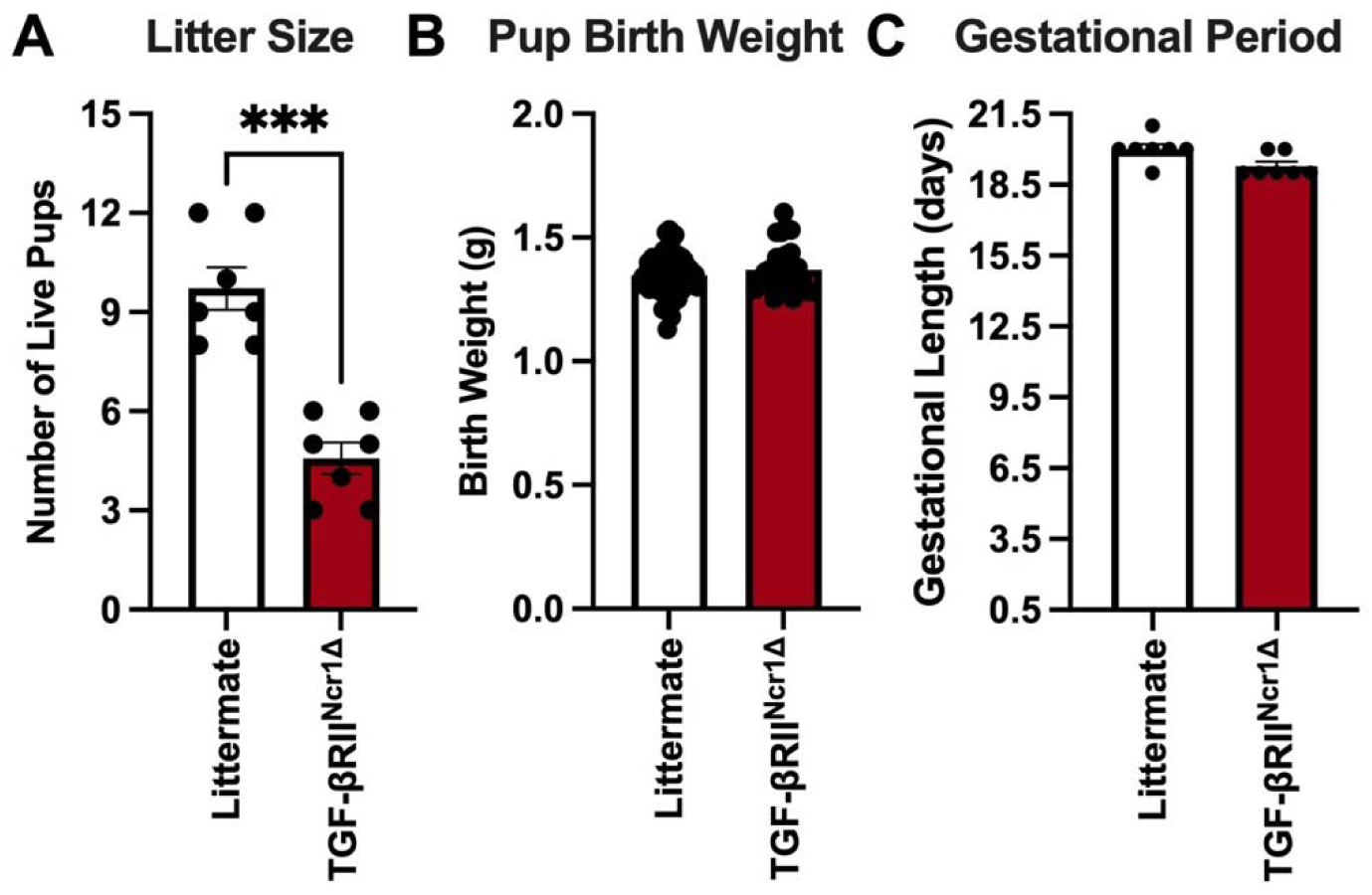
Impaired TGF-β–dependent uterine trNK cells differentiation leads to adverse pregnancy outcomes characterized by reduced litter sizes. (**A**) Number of live pups at first parturition from littermate control and TGF-βRII^Ncr1Δ^ dams (Littermates, *n*=7; TGF-βRII^Ncr1Δ^, *n*=7). (**B**) Pup birth weight in grams (g) from pups birthed by littermate control and TGF-βRII^Ncr1Δ^ dams (Littermates, *n*=68; TGF-βRII^Ncr1Δ^, *n*=31). (**C**) Gestational period in days for littermate control and TGF-βRII^Ncr1Δ^ dams (Littermates, *n*=7; TGF-βRII^Ncr1Δ^, *n*=7). Statistics were calculated using unpaired *t tests* with the Mann-Whitney correction. Error bars indicate SEM; *** *p* < 0.001.

uNK cells are thought to be critical regulators of placental vasculature remodeling^9,10,12,14^, particularly of the decidual spiral arteries, prompting us to investigate whether impaired uterine trNK cell differentiation due to loss of TGF-β signaling affects this process at midgestation. At gd 10.5, TGF-βRII^Ncr1Δ^ dams had fewer implantation sites and increased resorption rates compared to littermate controls (**Figure 4 A, B**). Furthermore, stereological quantification of decidual spiral arteries in midsagittal sections of gd 10.5 implantation sites revealed abnormalities in TGF-βRII^Ncr1Δ^ dams (**Figure 4 C**). Specifically, decidual spiral arteries from TGF-βRII^Ncr1Δ^ dams exhibited a 37% reduction in luminal area and a 75% increase in wall thickness, resulting in an increased vessel-to-lumen ratio relative to littermate controls (**Figure 4 D**). Taken together, these findings indicate that impaired uterine trNK cell differentiation in the absence of TGF-β signaling disrupts decidual spiral artery remodeling, leading to increased fetal resorptions at midgestation and an overall reduction in litter sizes at birth.

**Figure 4.**
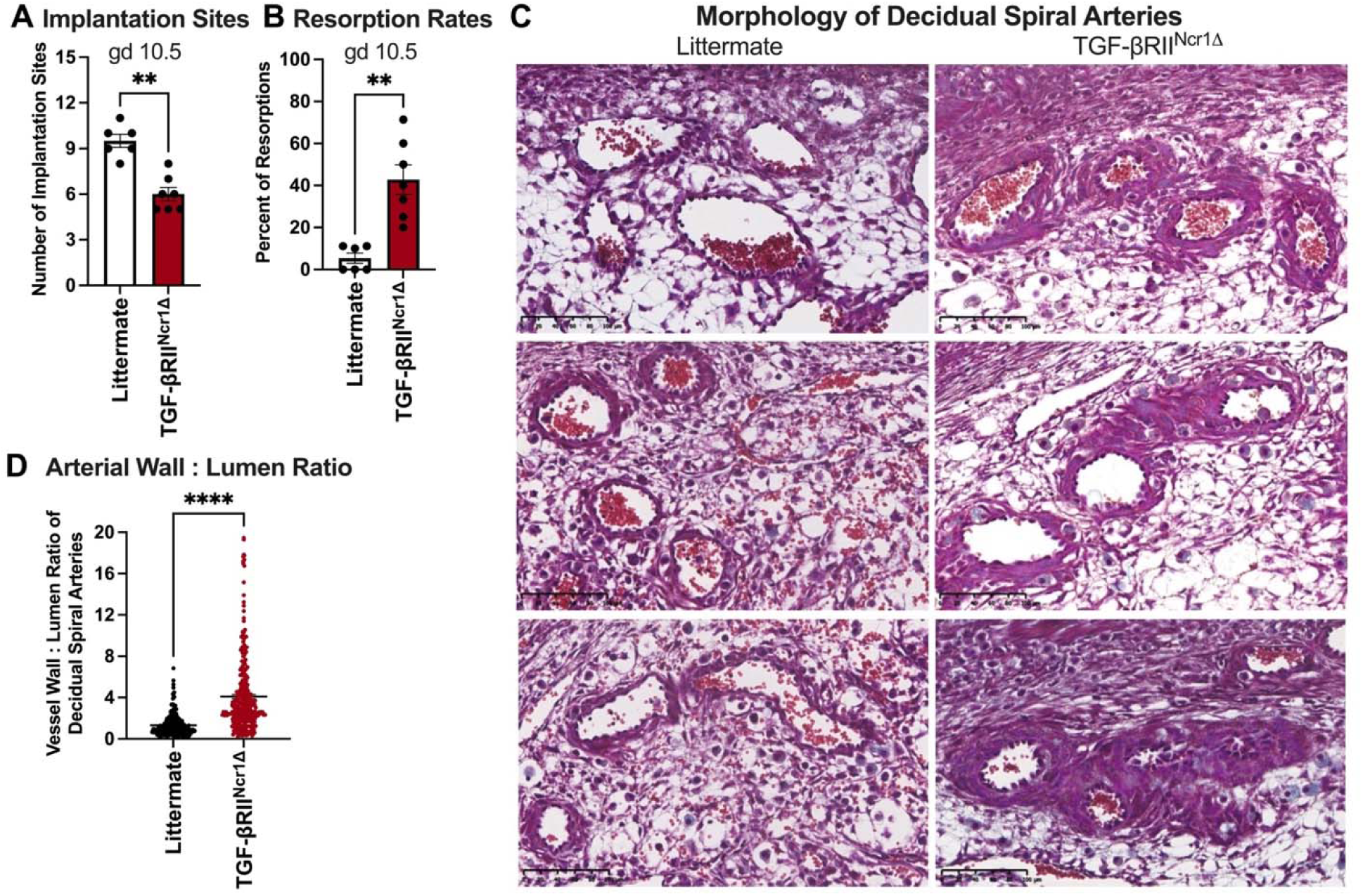
TGF-β–dependent uterine trNK cell differentiation required for proper spiral artery remodeling and fetal survival. (**A**) At gd 10.5, TGF-βRII^Ncr1Δ^ dams had fewer implantation sites than littermate control dams (Littermates, *n*=6; TGF-βRII^Ncr1Δ^, *n*=7). (**B**) Fetal resorption rates in littermate control and TGF-βRII^Ncr1Δ^ dams at gd 10.5, showing increased resorptions in conditional knockout dams (Littermates, *n*=6; TGF-βRII^Ncr1Δ^, *n*=7). Resorption rates (RR) were calculated as: RR(%) = (number of resorbed implantation sites/number of total implantation sites) X 100. (**C**) Representative images of gd 10.5 decidual spiral arteries from three littermate control and three TGF-βRII^Ncr1Δ^ dams stained with Masson’s Trichrome (Littermates, *n*=6; TGF-βRII^Ncr1Δ^, *n*=7; Scale bar, 100μm). (**D**) Spiral artery wall-to-lumen ratio at gd 10.5 implantation sites from littermate control and TGF-βRII^Ncr1Δ^ dams. Increased wall-to-lumen ratio in TGF-βRII^Ncr1Δ^ dams indicative of impaired spiral artery remodeling. (Littermates, *n*=6, decidual spiral arteries *n*=257; TGF-βRII^Ncr1Δ^, *n*=7 decidual spiral arteries *n*=305). Statistics were calculated unpaired *t tests* with the Mann-Whitney correction. Error bars indicate SEM; ** *p* < 0.01; and **** *p* < 0.0001.

### Partial reconstitution of uterine trNK cells restores midgestational pregnancy outcomes in TGF-βRII^Ncr1Δ^ dams

To determine whether restoring uterine trNK cells could rescue the midgestational pregnancy defects observed in TGF-βRII^Ncr1Δ^ dams, we adoptively transferred wildtype, congenically labeled splenic cNK cells into pregnant TGF-βRII^Ncr1Δ^ dams at gd 0.5. By gd 10.5, donor cNK cells were detected in the pregnant uterus, where a subset upregulated CD49a and downregulated CD49b, consistent with acquisition of a uterine trNK cell phenotype (**Figure 5 A**). However, adoptively transferred splenic cNK cells only partially reconstituted the uterine trNK cell population in the gravid uterus of TGF-βRII^Ncr1Δ^ dams, as evidenced by reduced absolute number and frequency of donor-derived trNK cells in reconstituted TGF-βRII^Ncr1Δ^ dams (**Figure 5 A-C**). Notably, this partial reconstitution was sufficient to rescue the gestational defects caused by impaired TGF-β–mediated uterine trNK cell differentiation. Reconstituted TGF-βRII^Ncr1Δ^ dams exhibited implantation site numbers and fetal resorption rates at gd 10.5 comparable to those observed in littermate controls (**Figure 5 D, E**). Together, these findings suggest that even partial restoration of the uterine trNK cell in pregnant TGF-βRII^Ncr1Δ^ dams is sufficient to restore pregnancy outcomes at midgestation, supporting a central role for uterine trNK cells as the principal NK cell subset required for successful murine pregnancy.

**Figure 5.**
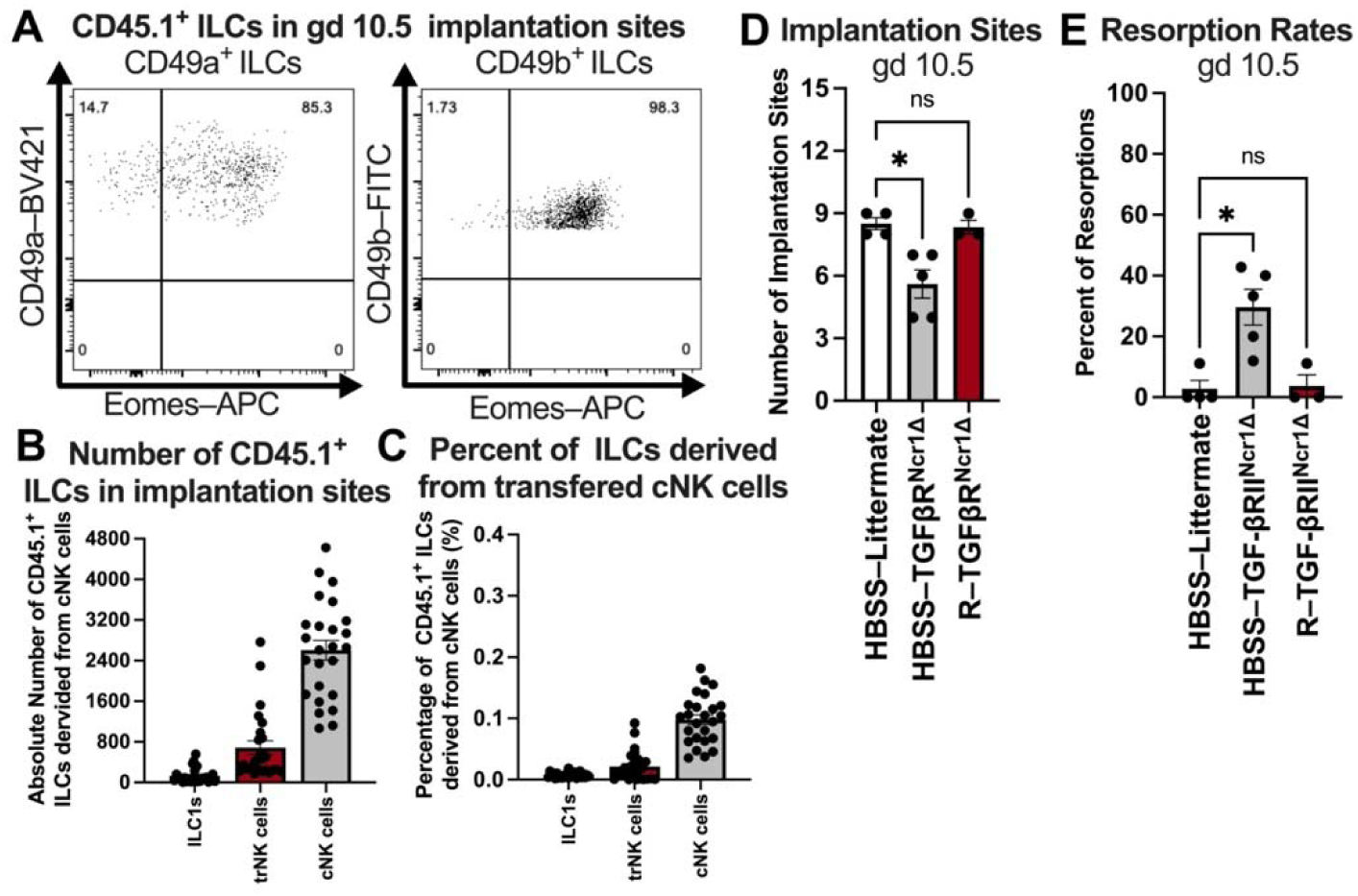
Adoptive transfer of splenic cNK cells partially reconstitutes uterine trNK cells and rescues midgestational pregnancy defects in TGFBRII^Ncr1Δ^ dams. (**A**) Representative flow plots showing the expression of CD49a, CD49b, and Eomes across CD45.1^+^ ILC subsets in gd 10.5 implantation sites of TGF-βRII^Ncr1Δ^ dams reconstituted with splenic CD45.1^+^ cNK cells. Briefly, 3.0x10^6^ splenic CD45.1^+^ cNK cells were adoptively transferred into TGF-βRII^Ncr1Δ^ dams at gd 0.5. By gd 10.5, a portion of adoptively transferred cNK cells in pregnant uterus of TGF-βRII^Ncr1Δ^ dams upregulated CD49a and downregulated CD49b, acquiring a CD49a^+^ CD49b^-^ Eomes^+^ phenotype characteristic of uterine trNK cells (Reconstituted (R)–TGF-βRII^Ncr1Δ^, *n*=3, implantation sites, *n*=25). Gating strategy: Live, Single Cells; CD3^-^ CD19^-^ CD45.1^+^ CD45.2^-^ NK1.1^+^ NKp46^+^ cells. (**B**) Absolute numbers of CD45.1^+^ ILC subsets in gd 10.5 implantation sites from reconstituted TGF-βRII^Ncr1Δ^ dams (R–TGF-βRII^Ncr1Δ^, *n*=3, implantation sites, *n*=25). (**C**) Proportion of uterine CD45.1^+^ ILC subsets derived from adoptively transferred splenic cNK cells in gd 10.5 implantation sites from TGF-βRII^Ncr1Δ^ dams (R–TGF-βRII^Ncr1Δ^, *n*=3, implantation sites *n*=25). (**D**) Number of implantation sites and (**E**) fetal resorption rates in reconstituted TGF-βRII^Ncr1Δ^ dams at gd 10.5 were comparable to those measured in littermate control dams injected intravascularly with HBBS (HBSS–Littermates, *n*=4; R–TGF-βRII^Ncr1Δ^, *n*=3). Resorption rates (RR) were calculated as: RR(%) = (number of resorbed implantation sites/number of total implantation sites) X 100. Statistics were calculated unpaired *t tests* with the Mann-Whitney correction.

## Discussion

Building on our previous work indicating that trNK cells in the pregnant uterus are derived from the cNK cell lineage, we now demonstrate that TGF-β signaling during murine gestation directs the differentiation of peripheral cNK cells into uterine trNK cells during murine gestation. This process is crucial for maintaining decidual spiral artery integrity and supporting successful pregnancy outcomes. This study represents the first direct *in vivo* evidence that TGF-β signaling mediates the generation of uterine trNK cells from peripheral cNK cells, uncovering an unrecognized role for TGF-β in shaping the distinct composition of innate lymphocytes at the maternal-fetal interface.

Our adoptive transfer studies demonstrate that uterine trNK cells can arise from a plastic reprogramming of peripheral cNK cells upon entry into the gravid uterus, highlighting the uterus as a dynamic and instructive niche that directs cNK cells to acquire a specialized tissue-specific phenotype. Consistent with our findings, recent human uterine transplantation studies demonstrate that certain subsets of uNK cells can be replenished from recipient-derived cells in the grafted uterus, supporting a model of continuous uNK cell differentiation^25^. Importantly, our studies suggest this phenotypic conversion is not a passive consequence of tissue-residency but is actively instructed by TGF-β signaling in the pregnant uterus. By identifying TGF-β as a central regulator directing uterine trNK cell differentiation during murine gestation, our work provides a mechanistic explanation of the process by which trNK cells acquire specialized tissue-specific phenotypes within the uterus that help sustain pregnancy, reconciling the long-standing paradox of why uterine trNK cells differ so markedly from their circulating counterparts.

The absence of cNK cell accumulation in the gravid uterus in the setting of impaired TGF-β signaling suggests a defect in tissue retention rather than recruitment. In the absence of TGF-β–mediated cues, circulating cNK cells that enter the uterine vasculature may fail to acquire the molecular programs required for residency and instead continue to transit through the tissue. This is consistent with a model in which TGF-β signaling promotes not only phenotypic conversion but also the acquisition of retention signals necessary for persistence within the uterine microenvironment, reinforcing that acquisition of tissue-residency in the gravid uterus is an actively instructed process.^29,32^

The functional consequences of this TGF-β–dependent pathway of uterine trNK cell differentiation on fecundity are profound. Impaired trNK cell differentiation in the absence of TGF-β signaling resulted in adverse pregnancy outcomes in mice, characterized by reduced litter sizes at first parturition. While our prior studies suggest uNK cells are critical for gestation, how these cells ensure pregnancy success remains poorly understood. In this study, impaired TGF-β–dependent trNK cell differentiation was associated with abnormal spiral artery remodeling and increased fetal resorption at midgestation. Notably, partial reconstitution of the uterine trNK cell compartment in TGF-βRII^Ncr1Δ^ dams was sufficient to normalize implantation site numbers and fetal resorption rates at midgestation. Together, these findings position uterine trNK cells as key contributors to placental vascular adaptations in the gravid uterus, emphasizing the importance of TGF-β–driven uterine trNK cell differentiation for murine reproductive success. The involvement of uNK cells in decidual spiral artery remodeling has been previously inferred from histological comparisons of implantation sites from immunodeficient mouse models; however, our work expands on these observations by pinpointing the subset of uNK cells involved in decidual angiogenesis^8–11^. Additional studies are necessary to elucidate the molecular mechanisms through which uterine trNK cells remodel decidual spiral arteries in the pregnant mouse uterus.

Interestingly, the inability to fully reconstitute the uterine trNK cell compartment following adoptive transfer suggests that only a subset of circulating cNK cells may be capable of differentiating into trNK cells during pregnancy, or alternatively that trNK cells already present in the virgin uterus may undergo *in situ* proliferation in the gravid uterus. Previous studies from our lab as well as others show that trNK cells within the pregnant murine uterus express marked levels of Ki67, supporting a model in which local proliferation of uterine trNK cells is a major contributor to the uterine trNK cell pool during pregnancy^7,33^. Prior studies have also described hematopoietic precursors within endometrial and decidual tissues that generate uterine trNK cells, suggesting that the compartment may be also sustained by local precursor differentiation^34–36^. Together, these findings suggest that uterine trNK cell ontogeny may be more complex than a single-source model and raise the possibility that distinct developmental pathways may operate at different stages of reproductive life. Therefore, defining the relative contribution and developmental timing of hematogenous versus locally maintained sources *in vivo* could provide relevant insights into the developmental trajectories and transcriptional programs that underlie decidual NK cell heterogeneity.

Notably, a reduction in the total number of implantation sites was also observed in TGF-βRII^Ncr1Δ^ dams, suggesting impaired TGF-β–dependent uterine trNK cell differentiation compromises implantation success. This finding raises the possibility that uterine trNK cells contribute to pregnancy not only through the remodeling of the placental vasculature but also by promoting uterine receptivity to implantation. In humans, studies of recurrent implantation failure and recurrent miscarriages have similarly implicated uNK cells in establishing endometrial receptivity and subsequently facilitating embryo implantation^22,23,37,38^. Therefore, future investigation is warranted to determine whether TGF-β–dependent alterations in uterine trNK cell–derived signals coordinate the cellular crosstalk underlying embryo implantation.

In addition to its effects on uterine trNK cells, loss of TGF-β receptor II in *Ncr1*-expressing cells substantially reduced ILC1s in the gravid murine uterus. However, the functional relevance of uterine ILC1s in pregnancy appears limited. Our previous findings show that the residual ILC1 population present in the gravid uterus of Eomes^Ncr1Δ^ dams failed to ameliorate the adverse pregnancy outcomes observed in this model, suggesting that uterine ILC1s are not required for successful gestation in mice^17^. Thus, the loss of ILC1s in the gravid uterus of TGF-βRII^Ncr1Δ^ dams is unlikely to account for the pregnancy defects detected in this mouse model.

More broadly, the role of TGF-β as a master regulator that tempers NK cell effector function extends beyond the pregnant uterus, shaping NK cell phenotype and functions across diverse tissue microenvironments. In the tumor microenvironment, TGF-β signaling suppresses NK cell toxicity by driving their conversion toward a ILC1-like phenotype that facilitates tumor immune evasion^30^. Similarly, in the obese murine liver, cNK cells undergo a TGF-β–dependent shift toward a less cytotoxic, ILC1-like state that mitigates tissue injury and protects against nonalcoholic fatty liver disease^31^. Within the gravid murine uterus, this same signaling axis is harnessed to promote maternal-fetal tolerance by reprograming peripheral cNK cells into uterine trNK cells with specialized, proangiogenic functions that sustain fetal development rather than immune activation. Whether a similar TGF-β driven program of uterine trNK cell differentiation is conserved in human pregnancy remains an important question for future investigation. Collectively, these findings position TGF-β as a context-dependent modulator of NK cell identity, fine-tuning their phenotype and function to the physiological needs of the surrounding microenvironment–whether to limit inflammation, permit tumor growth, or ensure reproductive success.

In conclusion, our work supports a model of continuous uterine trNK cell differentiation during murine gestation, in which peripheral cNK cells are recruited to the gravid uterus and converted into trNK cells via TGF-β signaling. This dynamic differentiation pathway ensures that uNK cells are appropriately tuned to the unique physiologic needs of gestation, linking NK cell plasticity directly to reproductive success. Our findings establish TGF-β–driven uterine trNK cell differentiation as a central axis for immune regulation in pregnancy and provide a framework that will be important for future studies exploring how perturbations in this pathway could underlie pregnancy complications. These insights reshape our understanding of NK cell developmental plasticity and highlight uterine immune adaptation as a fundamental component of reproductive fitness.

## Materials and Methods

### Mice

All mouse studies were performed in accordance with ethical guidelines and animal protocol approved by the Washington University School of Medicine Animal Studies Committee under protocol number *24-0332.* Wild-type *C57BL/6* (stock number 665) and *B6.SJL-Ptprc^a^Pecpc^b^/BoyJ* (stock number 664) mice were purchased from Charles River Laboratories (Wilmington, MA). *Ncr1^iCre^* were generously gifted by Eric Vivier at Aix Marseille University, Marseille, France. TGF-βRII^Ncr1Δ^ mice and littermate controls were generated by crossing *B6;129-Tgfbr2^tm1Karl^/J* (strain: 012603; The Jackson Laboratory, Bar Harbor, ME) with *Ncr1^iCre^*mice. Female mice aged 6-8 weeks were used in all mouse studies. All mice were housed in the Laboratory for the Animal Care barrier facility at the Washington University School of Medicine and maintained on a 12-hour light/dark cycle.

### Timed-Pregnancies

To distinguish maternal lymphocytes from fetal lymphocytes at the maternal-fetal interface, we mated virgin C57BL/6, littermate control, Ncr1^iCre^ and TGF-βRII^Ncr1Δ^ female mice with C57BL/6 CD45.1 male mice overnight. The timing of conception was determined by detection of a copulation plug the following morning, which was designated as gd 0.5. Pregnancy outcomes were evaluated by comparing gestational length, litter sizes, and pup birth weight at first parturition between littermate control and TGF-βRII^Ncr1Δ^ dams. Pregnant dams were dissected at gds 6.5, 10.5, or 14.5 to assess the immune constituents of implantation sites, and at gd 10.5 to assess fetal resorption rates and placental vasculature morphology. Fetal resorption rates were calculated as the percentage of resorbed implantation sites per pregnant uterus.

### cNK Cell Adoptive Transfer Studies in Pregnant Dams

Wildtype CD45.2 splenic cNK cells were purified by negative selection (Stem Cell Technologies, Vancouver, Canada) to obtain CD3^-^ CD19^-^ NK1.1^+^ NKp46^+^ CD49b^+^ cNK cells with purity of 87-93%. 2.5x10^6^ purified CD45.2^+^ cNK cells were injected intravascularly via tail vein injection into C57BL/6 CD45.1 pregnant dams at gd 0.5. The phenotype of adoptively transferred cNK cells was then assessed by flow cytometry at gd 10.5.

### cNK Cell Adoptive Transfer Studies in Pregnant TGF-βRII^Ncr1Δ^ Dams

Wildtype CD45.1^+^ splenic cNK cells were purified by negative selection (Stem Cell Technologies, Vancouver, Canada) to obtain CD3^-^ CD19^-^ NK1.1^+^ NKp46^+^ CD49b^+^ cNK cells with purity of 85-92%. 3.0x10^6^ purified CD45.1^+^ cNK cells were suspended in 300 μl of sterile Hank’s Balanced Salt Solution (HBSS) and injected intravascularly via tail vein injection into pregnant TGF-βRII^Ncr1Δ^ dams at gd 0.5. Littermate control dams were injected with 300 μl of sterile HBSS intravascularly via tail vein injection. The phenotype of adoptively transferred cNK cells in reconstituted TGF-βRII^Ncr1Δ^ dams was then assessed by flow cytometry at gd 10.5. The number of implantation sites and fetal resorption rates were assessed at gd 10.5. as described above in reconstituted TGF-βRII^Ncr1Δ^ dams and HBSS-treated littermate control dams.

### Intravascular Staining of Circulating Lymphocytes

To distinguish tissue-resident lymphocytes at the maternal-fetal interface from those in circulation, we administered 3 μg of fluorescently labeled anti-CD45.2–PE-Cy7 (104; Invitrogen, Waltham, MA) to pregnant dams intravascularly via tail vein injection 3 minutes prior to euthanasia.

### Single Cell Isolations from Different Tissues

#### Implantation Site Digestion

Pregnant dams were dissected at gds 6.5, 10.5, or 14.5 to assess the immune constituents of individual implantation sites. Each healthy implantation was digested with Liberase TL (167 μg/ml; Sigma-Aldrich, St. Louis, MO) and DNase1 (150 μg/ml; Sigma-Aldrich, St. Louis, MO) for 1 hour at 37°C. Enzymatically digested implantation sites were minced, washed with 10% FBS RPMI media, and resuspended in 3mL of complete R10 media.

#### Splenic Preparation

Spleens from each pregnant dam were harvested, minced, and filtered through a 70 μm mesh. Splenocyte suspensions were subsequently treated with RBC Lysis Buffer, washed with 10% FBS RPMI media, and resuspended in 5mL of complete R10 media.

### Flow Cytometry

Fluorescently-labeled antibodies to the indicated antigens were purchased from the following vendors: Invitrogen (Waltham, MA), which included CD3e (clone 145-2C11), CD19 (1D3), CD45.2 (104), CD45.1 (A20), CD49b (DX5), EOMES (Dan11mag), NKp46 (29A1.4), and Fixable Viability Dye (eFluorTM506); BioLegend (San Diego, CA) which included NK1.1 (PK136); BD Biosciences (Franklin Lakes, NJ) which included CD49a (Ha31/8) and Streptavidin (PE-Conjugate); R&D Systems (Minneapolis, MN) which included TGF-βRII (Biotinylated).

Prior to staining, 5,000 Precision Count Beads (BioLegend, San Diego, CA) were added to each sample to quantify cell numbers. Cells were stained with fixable viability dye (Invitrogen, Waltham, MA) and then stained for cell surface markers in 2.4G2 hybridoma supernatant (anti-FcγRIII) to block Fc receptors. Following surface staining, cells were fixed and permeabilized using the Foxp3/Transcription Factor Staining Buffer Set (eBiosciences; San Diego, CA) according to the manufacturer’s instructions and subsequently stained for intracellular molecules. All samples were acquired on a FACS Canto (BD Biosciences, Franklin Lakes, NJ) and analyzed using FlowJo Software 10.8.2 (BD Biosciences, Franklin Lakes, NJ). Maternal uterine cNK cells were defined as viable singlets, CD3^-^ CD19^-^ CD45.1^-^ CD45.2^+^ NK1.1^+^ NKp46^+^ CD49a^-^ CD49b^+^ Eomes^+^. Maternal uterine trNK cells were defined as viable singlets, CD3^-^ CD19^-^CD45.1^-^ CD45.2^+^ NK1.1^+^ NKp46^+^ CD49a^+^ CD49b^-^ Eomes^+^. Maternal uterine ILC1s were defined as viable singlets CD3^-^ CD19^-^ CD45.1^-^ CD45.2^+^ NK1.1^+^ NKp46^+^ CD49a^+^ CD49b^-^ Eomes^-^.

### Morphological Analysis of Decidual Spiral Arteries

Placental tissues at gd 10.5 were prepared for histology by fixing uterine horns containing intact implantation sites in 4% paraformaldehyde, followed by paraffin embedding. Thin tissue sections (5 μm) were cut and processed for Masson’s Trichrome staining according to manufacturer’s instructions. All slides were examined by light microscopy, and images were captured using a NanoZoomer HT2.0 Digital Slide Scanner (Hamamatsu Photonics, Hamamatsu, Japan). The center section of each serially sectioned implantation site was identified, and at least 4 implantation sites per litter were analyzed for each dam. Vessel wall and lumen measurements of decidual spiral arteries were taken from cross-sectional images using NIS-Elements Viewer NDP.view2 4.11.0 Imaging Software (Nikon Microscope). Vessel wall-to-lumen ratios for individual decidual spiral arteries were calculated as the ratio of the outer wall area to the luminal area. Investigators were blinded to sample genotypes during analysis.

### Statistical Analysis

Statistical analysis was performed with Prism 10.4.1 (GraphPad software) using unpaired *t tests* with the Mann-Whitney correction. Error bars in figures represent the SEM. Normality was assessed with Prism 10.4.1 (GraphPad software) using Q-Q plots. Sample sizes for each experiment were determined using a significance level (α) of 0.5 and power of 85%. Statistical significance was indicated as follows: ns, not significant; \**p* < 0.05; ** *p* < 0.01; *** *p* < 0.001; and **** *p* < 0.0001.

## Acknowledgements

We extend our gratitude to the members of the Yokoyama Laboratory for their insightful discussions and constructive feedback. Additionally, we thank the Alafi Neuroimaging Core for access to the NanoZoomer HT2.0 Digital Slide Scanner, supported by the shared instrumentation grant NCRR1S10RR027552. This work was supported by the Eunice Kennedy Shriver National Institute of Child Health and Human Development of the National Institutes of Health under award number 1F30HD118750-01A1.

## Abbreviations used in this paper include

cNK: conventional NK
Eomes: Eomesodermin
Gd: gestational day
ILC: innate lymphoid cell
ILC1: type 1 innate lymphoid cell
TGF-β: transforming growth factor-β
trNK: tissue-resident NK
uNK: uterine NK

## Data Availability

Data supporting the findings of this study are available from the corresponding author upon reasonable request.

## Authors’ Contributions

J.D.B., performed experiments; acquisition and analysis of data; drafting of the manuscript; J.D.B., D.M.N., and W.M.Y, study concept and design; interpretation of data; critical revision of the manuscript; L.Y., technical support; W.M.Y., provided supervision; obtained funding. All authors approved of the final version of this manuscript.

---

**Supplementary Figure 1.**
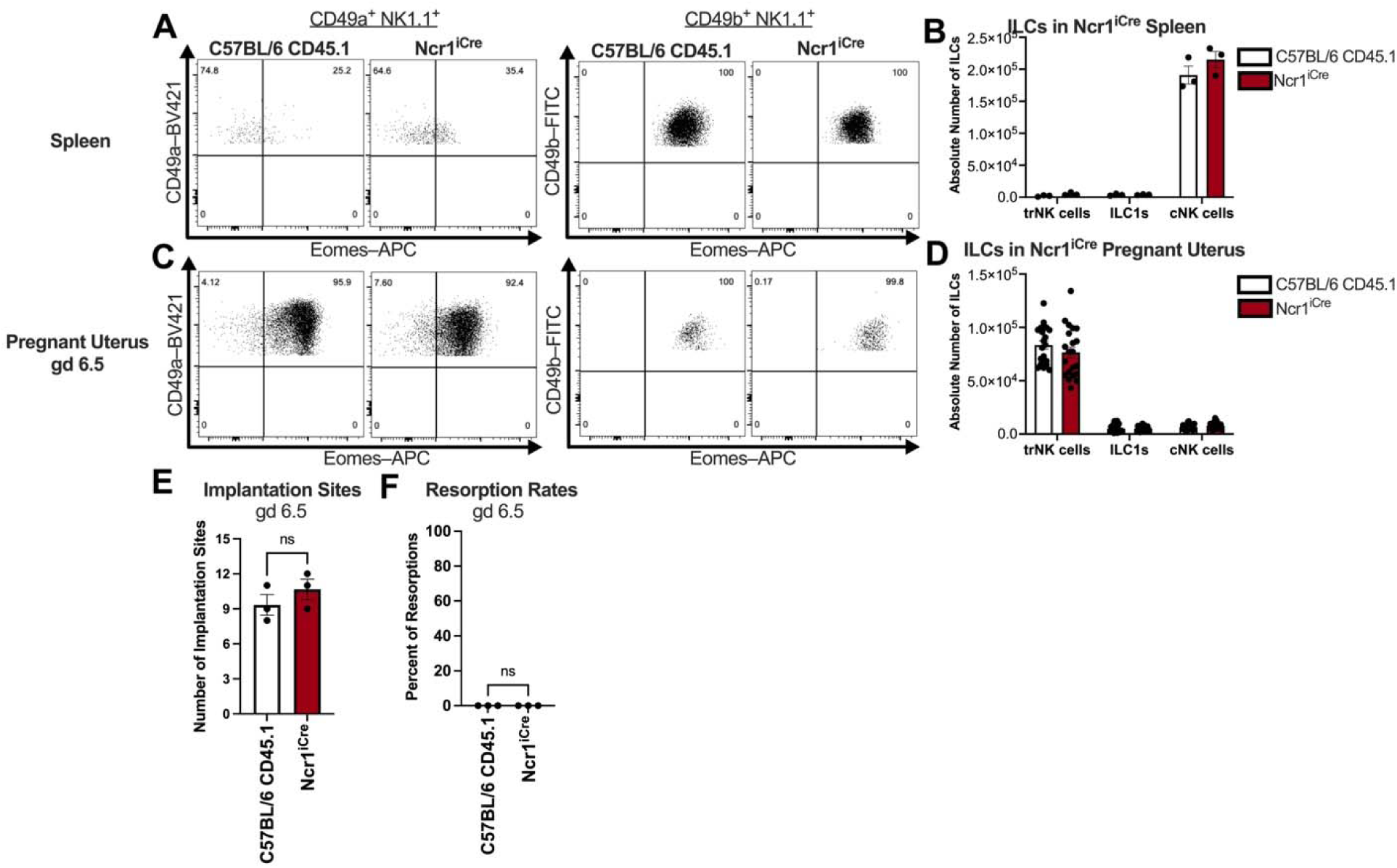
Ncr1^iCre^ expression does not alter uterine ILC composition or early pregnancy outcomes. (**A**) Representative flow plots showing the expression of CD49a, CD49b, and Eomes across ILC subsets in the spleens of wildtype C57BL/6 CD45.1 and Ncr1^iCre^ dams at gd 6.5 (C57BL/6 CD45.1, *n*=2; Ncr1^iCre^*, n*=3). Gating strategy: Live, Single Cells; CD3^-^ CD19^-^ CD45.1^-^ CD45.2^+^ NK1.1^+^ NKp46^+^ cells. (**B**) Absolute cell counts of CD49a^+^ Eomes^+^ trNK cells, CD49a^+^ Eomes^-^ ILC1s, and CD49b^+^ Eomes^+^ cNK cells in the spleens of pregnant wildtype C57BL/6 CD45.1 and Ncr1^iCre^ dams at gd 6.5. (**C**) Representative flow plots showing the expression of CD49a, CD49b, and Eomes across ILC subsets in the gravid uterus of wildtype C57BL/6 CD45.1 and Ncr1^iCre^ dams at gd 6.5 (C57BL/6 CD45.1, *n*=3, implantation sites *n*=21; Ncr1^iCre^, *n*=3, implantation sites *n*=21). (**D**) Absolute cell counts of CD49a^+^ Eomes^+^ trNK cells, CD49a^+^ Eomes^-^ ILC1s, and CD49b^+^ Eomes^+^ cNK cells in the gravid uterus of wildtype C57BL/6 CD45.1 and Ncr1^iCre^ dams at gd 6.5. (**E**) Number of implantation sites and (**F**) fetal resorption rates in Ncr1^iCre^ dams at gd 6.5 were comparable to those measured in wildtype C57BL/6 CD45.1 dams (C57BL/6 CD45., *n*=3; Ncr1^iCre^ dams, *n*=3). Resorption rates (RR) were calculated as: RR(%) = (number of resorbed implantation sites/number of total implantation sites) X 100. Statistics were calculated unpaired *t tests* with the Mann-Whitney correction.

